# The distribution and evolutionary dynamics of dopaminergic neurons in molluscs

**DOI:** 10.1101/2024.06.26.600886

**Authors:** Tigran P. Norekian, Leonid L. Moroz

**Author notes:** **Corresponding author:** Dr. Leonid L. Moroz.

## Abstract

Dopamine is one of the most versatile neurotransmitters in invertebrates. It’s distribution and plethora of functions is likely coupled to feeding ecology, especially in Euthyneura (the largest clade of molluscs), which presents the broadest spectrum of environmental adaptations. Still, the analyses of dopamine-mediated signaling were dominated by studies of grazers. Here, we characterize the distribution of dopaminergic neurons in representatives of two distinct ecological groups: the sea angel - obligate predatory pelagic mollusc *Clione limacina* (Pteropoda, Gymnosomata) and its prey - the sea devil *Limacina helicina* (Pteropoda, Thecosomata) as well as the plankton eater *Melibe leonina* (Nudipleura, Nudibranchia). By using tyrosine hydroxylase-immunoreactivity (TH-ir) as a reporter, we showed that the dopaminergic system is moderately conservative among euthyneurans. Across all studied species, small numbers of dopaminergic neurons in the central ganglia contrast to significant diversification of TH-ir neurons in the peripheral nervous system, primarily representing sensory-like cells, which predominantly concentrated in the chemotactic areas and projecting afferent axons to the central nervous system. Combined with α-tubulin immunoreactivity, this study illuminates the unprecedented complexity of peripheral neural systems in gastropod molluscs, with lineage-specific diversification of sensory and modulatory functions.

## 1 INTRODUCTION

As a neuronal messenger, 3,4-dihydroxy<u>p</u>henethylamine (3-hydroxytyramine or dopamine) was identified in 1957 (Carlsson et al., 1957; Montagu, 1957; Carlsson et al., 1958) together with localization of catecholaminergic neurons in the brain (Carlsson et al., 1962) by aldehyde-induced fluorescence techniques (Falck, 1962; Falck et al., 1962). Dopamine (DA) is one of the well-known and still one of the most mysterious neurotransmitters in molluscs and invertebrates in general. As with many monoamines, DA apparently was co-opted into neural functions in the common ancestor of bilaterians (Moroz et al., 2021) with a broad distribution across phyla. However, the generalized theme for DA and other catecholamines’ distribution and functions among invertebrates is still enigmatic (Walker et al., 1996; Spencer et al., 1998; Yamamoto and Vernier, 2011a; Gallo et al., 2016; Bauknecht and Jékely, 2017), primarily due to small sizes of dopaminergic neurons and a plethora of modulatory roles, including complex contribution to learning, memory and reward mechanisms even in well-studied model systems (Hollerman and Schultz, 1998; Schultz, 1998; Brembs et al., 2002; Hills et al., 2004; Andretic et al., 2005; Kume et al., 2005; Barron et al., 2010; Rivard et al., 2010; Waddell, 2010; Yamamoto and Vernier, 2011b; Fabbri et al., 2024; McMillen and Chew, 2024).

Gastropods of the highly diverse clade Euthyneura, subclass Heterobranchia, represent ∼40% (or ∼30,000) of the described molluscan species (Rosenberg, 2014; Bouchet et al., 2017; Brenzinger et al., 2021). Euthyneurans were particularly impactful in deciphering dopaminergic systems (Miller, 2020) because of the unique features of their central nervous systems, with multiple large visually identified neurons located at the ganglionic surface (Gillette, 1991), facilitating deciphering cellular bases of behaviors (Kandel, 1976; Chase, 2002) as well as neuroplasticity mechanisms (Kandel, 2001).

In these species, DA functions were associated with the interneuronal control of feeding (Rosen et al., 1991; Teyke et al., 1993; Kabotyanski et al., 1998; Díaz-Ríos and Miller, 2005; Neveu et al., 2017; Miller, 2020) and respiratory (Budko and Moroz, 1988; Moroz, 1990; Syed et al., 1990; Moroz, 1991; Syed and Winlow, 1991; Moroz and Winlow, 1992a; Inoue et al., 2001) circuits, valuation of sensory information, and reward reinforcements (Scheibenstock et al., 2002; Miller, 2020; Momohara et al., 2022).

The distribution of DA-containing neurons was the primary focus of multiple studies, including the identification of an asymmetric giant dopaminergic neuron in the pedal ganglia of freshwater pulmonate molluscs (Pentreath et al., 1974; Winlow et al., 1981) and relatively small number of buccal and cerebral dopaminergic neurons with moderate conservation across species (reviewed by (Miller, 2020).

Surprising funding was the evidence that many DA-containing neurons were not located in the CNS but at the periphery (Salimova et al., 1987; Croll, 2001; Martínez-Rubio et al., 2009; Horváth et al., 2020) emphasizing putative sensory signaling such as mechano- and chemoreception (Cummins et al., 2009), and those contributing to the activation of feeding motor outputs (Kabotyanski et al., 1998; Brown et al., 2018; Norekian et al., 2024).

The diversity of DA-containing cell populations both in the CNS and at the periphery among molluscs was linked to the feeding and, apparently, respiratory ecology, with euthyneurans representing the broadest spectrum of environmental adaptations and food consumption strategies.

Still, the analyses of DA-mediated signaling were dominated by studies of grazers such as *Aplysia* (Ascher et al., 1967; Swann et al., 1978; Croll, 2001; Díaz-Ríos et al., 2002), *Lymnaea* (Cottrell et al., 1979; Audesirk, 1985; Croll and Chiasson, 1990; Elekes et al., 1991; Elekes, 1992; Winlow et al., 1992; Magoski et al., 1995; Croll et al., 1999; Voronezhskaya et al., 1999; Young et al., 2022), *Biomphalaria* (Chiang et al., 1974; Vallejo et al., 2014; Vaasjo et al., 2018), *Helisoma* (Trimble et al., 1984; Harris and Cottrell, 1995; Quinlan et al., 1997; Kiehn et al., 2001) and *Helix* (Hernádi et al., 1993; Ierusalimsky et al., 1997) – the most popular reference species (Wentzell et al., 2009; Miller, 2020), which are relatively easy to maintain in the laboratory. In contrast, the carnivorous molluscs received less attention (Brown et al., 2018; Norekian et al., 2024), and it is understandable due to the methodological difficulties of finding and working with these species.

Here, we extend comparative aspects of the distribution of catecholaminergic neurons targeting two distinct ecological and less investigated groups of Euthyneura: the sea angel - obligate predatory pelagic mollusc *Clione limacina* (Pteropoda, Gymnosomata) and its prey - the sea devil -*Limacina helicina* (Pteropoda, Thecosomata) as well as the plankton eater *Melibe leonina* (Nudipleura, Nudibranchia, (Watson et al., 2021)), see **Fig. 1**. Pteropod molluscs are the sister lineage to *Aplysia* (Anaspidea), while Nudipleura represents one of the basal groups of Euthyneura (Dinapoli and Klussmann-Kolb, 2010; Bouchet et al., 2017), which justify the proposed selection of these species for comparative analyses.

**Figure 1.**
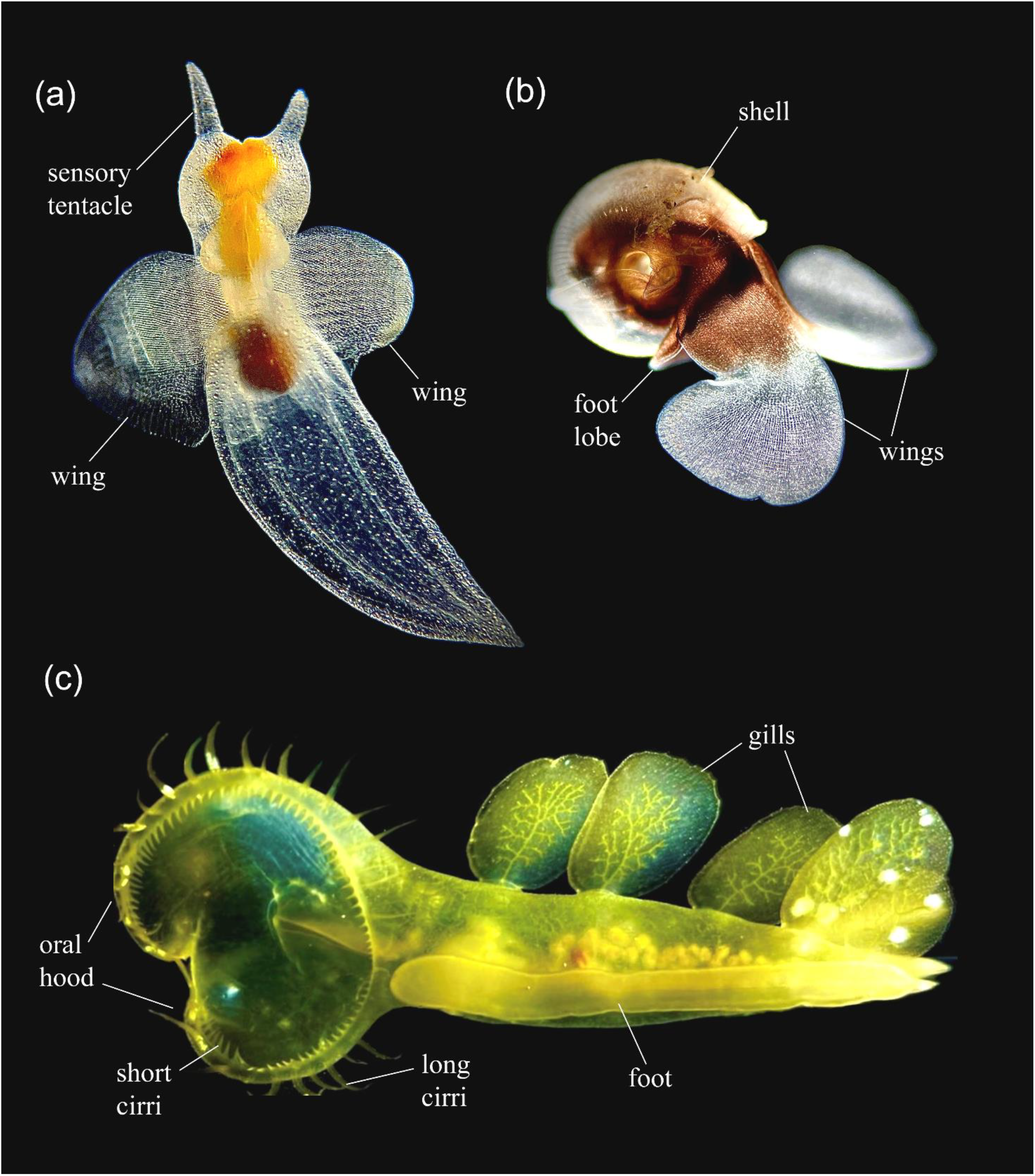
Images of live pteropod molluscs, *Clione limacina* (a), *Limacina helicina* (b) and the nudibranch *Melibe leonina* (c).

Both *Clione* and *Melibe* are recognized reference species in evolutionary neurobiology, with the identification of multiple neurons controlling feeding, swimming, and other behaviors (Sakharov, 1970; Gerasimov, 1973; Sakharov, 1974; Arshavsky Yu et al., 1985; Satterlie, 1985; Arshavsky Yu et al., 1989; Norekian, 1990a; Kabotyanskii and Sakharov, 1991; Page, 1992b; a; Satterlie, 1993; Norekian and Satterlie, 1996; Arshavsky et al., 1998; Deliagina et al., 1998; Moroz et al., 2000; Sadreyev and Panchin, 2000; Newcomb and Watson, 2001; 2002; Thompson and Watson, 2005; Kempf, 2008; Malyshev and Balaban, 2011; Satterlie, 2013; Duback et al., 2018; Sakurai and Katz, 2019; Pirtle, 2022). Nonetheless, very little is known about dopaminergic signaling (Sakharov and Kabotyanski, 1986; Kabotyanski and Sakharov, 1988; Norekian, 1990b) in these ecologically important groups, and the initial mapping of catecholaminergic neurons in *Clione* was performed using modified aldehyde-induced fluorescence techniques (Kabotyanski and Sakharov, 1989).

In this study, we used immunohistochemistry to localize tyrosine hydroxylase (TH), a specific enzyme in the pathway of catecholamine synthesis, which converts tyrosine into L-DOPA (Osborne et al., 1975; Osborne et al., 1976). Since dopamine is the only catecholamine found in significant quantities in gastropods (Carpenter et al., 1971; McCaman et al., 1979; Vallejo et al., 2014), TH-immunoreactivity serves as a good marker for dopaminergic neurons in gastropod molluscs, and a complementary method to map these cells and neuronal processes. It has been successfully used to identify dopaminergic neurons in the central ganglia and peripheral tissues of such gastropod species as *Pleurobranchaea californica, Aplysia californica, Helix pomatia, Helisoma*, and *Biomphalaria* species (reviewed by (Miller, 2020).

We characterized the distribution of the TH-immunoreactive (TH-ir) neural elements in *Clione, Limacina*, and *Melibe*, with a comparative analysis of the dopaminergic central and peripheral neural systems among them and other previously researched gastropods. We showed that the dopaminergic system is moderately conservative among euthyneurans but with varying numbers of neurons and ganglia. These observations revealed trends in the adaptive diversification of homologous neurons of different transmitter specificity. Across all studied species, small numbers of dopaminergic neurons in the central ganglia contrast to significant diversification of TH-ir neurons in the peripheral nervous system, primarily representing sensory-like cells, which predominantly concentrated in the chemotactic areas and projecting afferent axons to the central nervous system.

## 2 MATERIALS AND METHODS

### 2.1 Animals

Adult specimens of *Clione limacina, Limacina helicina*, and *Melibe leonina* (**Fig. 1**) were collected from the breakwater in the Northwest Pacific and held in 1-gal glass jars in large tanks with constantly circulating seawater at 10°C–12°C. Experiments were carried out at Friday Harbor Laboratories, the University of Washington, in the spring, summer, and fall seasons of 2020–2023.

### 2.2 Immunocytochemistry and Phalloidin staining

Animals were anesthetized in a 1:1 mixture of seawater and 0.3 M MgCl_2_ and dissected in Sylgard-coated Petri dishes. Adult *Clione* (1.5 to 2.5 cm long) had their body-tail separated, while the head and the wings remained connected to the central ganglia via head nerves and wing nerves. The buccal mass with the attached buccal ganglia was also separated. Smaller *Limacina* (about 1 cm in size) had their shell with internal organs removed, while the rest of the body with the wings and foot lobes was cut in two parts. Larger *Melibe* (5 to 12 cm long) had their central ganglia isolated; rhinophores cut off from the head, and edges of the feeding hood with sensory protrusions cut into several smaller manageable pieces. Prior to fixation, all isolated fragments of tissue were treated with a 1 mg/ml solution of protease (Sigma type XIV) for approximately 5 min, followed by a 20-30 min wash in seawater. This treatment helped to improve antibody penetration and remove connective tissues covering the central ganglia.

The dissected tissues were fixed overnight (10-12 hours) in 4% paraformaldehyde in 0.1 M phosphate-buffered saline (PBS) at +5º C and washed for 2 hours in PBS. The tissues were pre-incubated overnight in a 6% goat serum-blocking solution in PBS containing 0.02% Triton X-100 (PBT). To detect catecholaminergic neural elements, a mouse monoclonal antibody against tyrosine hydroxylase (the enzyme responsible for catalyzing the conversion of the amino acid L-tyrosine to L-DOPA, which is a precursor to dopamine) was used (Immunostar; Cat# 22941, RRID: AB_572268). Primary antibody dilution was 1:60 with an incubation time of 48 hours at 4°C. Following primary antibody incubation, the tissue was washed in PBS and incubated in secondary antibodies - goat anti-mouse IgG (Alexa 488 conjugated; Molecular Probes; Cat# A-11001, RRID: AB_2534069) at a final dilution of 1:30.

To label a wider group of neural elements in the peripheral tissue of all studied molluscs, we used the rat monoclonal antibody against tubulin (AbD Serotec Cat# MCA77G, RRID: AB_325003), which recognizes the alpha subunit of tubulin and binds explicitly tyrosylated tubulin (Wehland and Willingham, 1983; Wehland et al., 1983). Following a series of PBS washes for 6-8 hours, the dissected tissues were incubated for 24 hours in secondary goat anti-rat IgG antibodies: Alexa Fluor 488 conjugated (Molecular Probes, Invitrogen, Cat# A11006, RRID: AB_141373) at a final dilution 1:40.

We used the well-known marker phalloidin (Alexa Fluor 568 phalloidin from Molecular Probes) to label the muscle fibers, which binds to F-actin (Wulf et al., 1979). After washing in PBS following the secondary antibody treatment, the samples were incubated in phalloidin solution (in PBS) for 6 to 8 hours at +5º C, at a final dilution 1:80, and then washed in several PBS rinses for 8 hours.

To label cell nuclei, the tissue was mounted in Antifade Mounting Medium with DAPI (Vectashield; Cat#H-2000). Some of the larger adult tissues were also mounted in Fluorescent Mounting Media (KPL) on glass microscope slides. The slides were viewed using a Nikon Research Microscope Eclipse E800 with Epi-fluorescence using standard TRITC and FITC filters and recorded using Nikon C1 Laser Scanning confocal microscope.

### 2.2 Antibody specificity

Rat monoclonal antityrosinated alpha-tubulin antibody (Serotec Cat #MCA77G; RRID: AB_325003) has been extensively used in different species. As reported by Wehland et al. (1983) this rat monoclonal antibody “reacts specifically with the tyrosylated form of brain alpha-tubulin from different species” (Wehland and Willingham, 1983; Wehland et al., 1983). We have also successfully used this specific anti-α-tubulin antibody before on several species of ctenophores (Norekian and Moroz, 2016; 2020b) and hydrozoans (Norekian and Meech, 2020; Norekian and Moroz, 2020a) to label their nervous system. We also tested the specificity of immunostaining by omitting either the primary or the secondary antibody from the procedure. In both cases, no labeling was detected.

Mouse monoclonal antibody against tyrosine hydroxylase (Immunostar; Cat# 22941, RRID: AB_572268) has been previously used with success on the molluscs; for example, *Pleurobranchaea californica* (Brown et al., 2018; Norekian et al., 2024), *Aplysia californica* (Croll, 2001), *Helix pomatia* (Hernádi et al., 1993), *Lymnaea* (Elekes et al., 1991; Elekes, 1992), *Helisoma* (Kiehn et al., 2001) and *Biomphalaria* species (Vallejo et al., 2014). We also tested the specificity of immunostaining by omitting either the primary or the secondary antibody from the procedure. In both cases, no labeling was detected.

## 3 RESULTS

We selected three species of marine gastropod molluscs: (a) the gymnosome pteropod *Clione limacina*, without a shell; (b) the thecosome shelled pteropod *Limacina helicina*; and (c) hooded nudibranch *Melibe leonina* (**Fig. 1**). *Clione* and *Limacina* are both representatives of pelagic actively swimming pteropods, which life histories are mutually connected in oceanic ecosystems. *Clione* is a highly specialized predator that feeds only on *Limacina* (Wagner, 1885; Lalli and Gilmer, 1989). Because of such an exclusive predator-prey relationship, *Clione* has specialized feeding structures adapted for prey capture - 3 pairs of oral appendages, called buccal cones, that surround the mouth of *Clione* and, when retracted, are cone-shaped and covered by oral skin folds. **Fig. 2** shows that contact with the prey induces fast protraction of the buccal cones in *Clione*, which then become tentacle-like and seize the shell of the prey *Limacina* (Hermans and Satterlie, 1992; Norekian, 1993; Norekian and Satterlie, 1993a; b). The nudibranch mollusc *Melibe leonina* has a unique feeding behavior. It lacks radula and buccal mass, and uses movements of its large oral hood to capture food and bring it to the mouth to be swallowed without chewing into the stomach (Agersborg, 1921; Ajeska and Nybakken, 1976; Trimarchi and Watson, 1992; Watson and Trimarchi, 1992).

**Figure 2.**
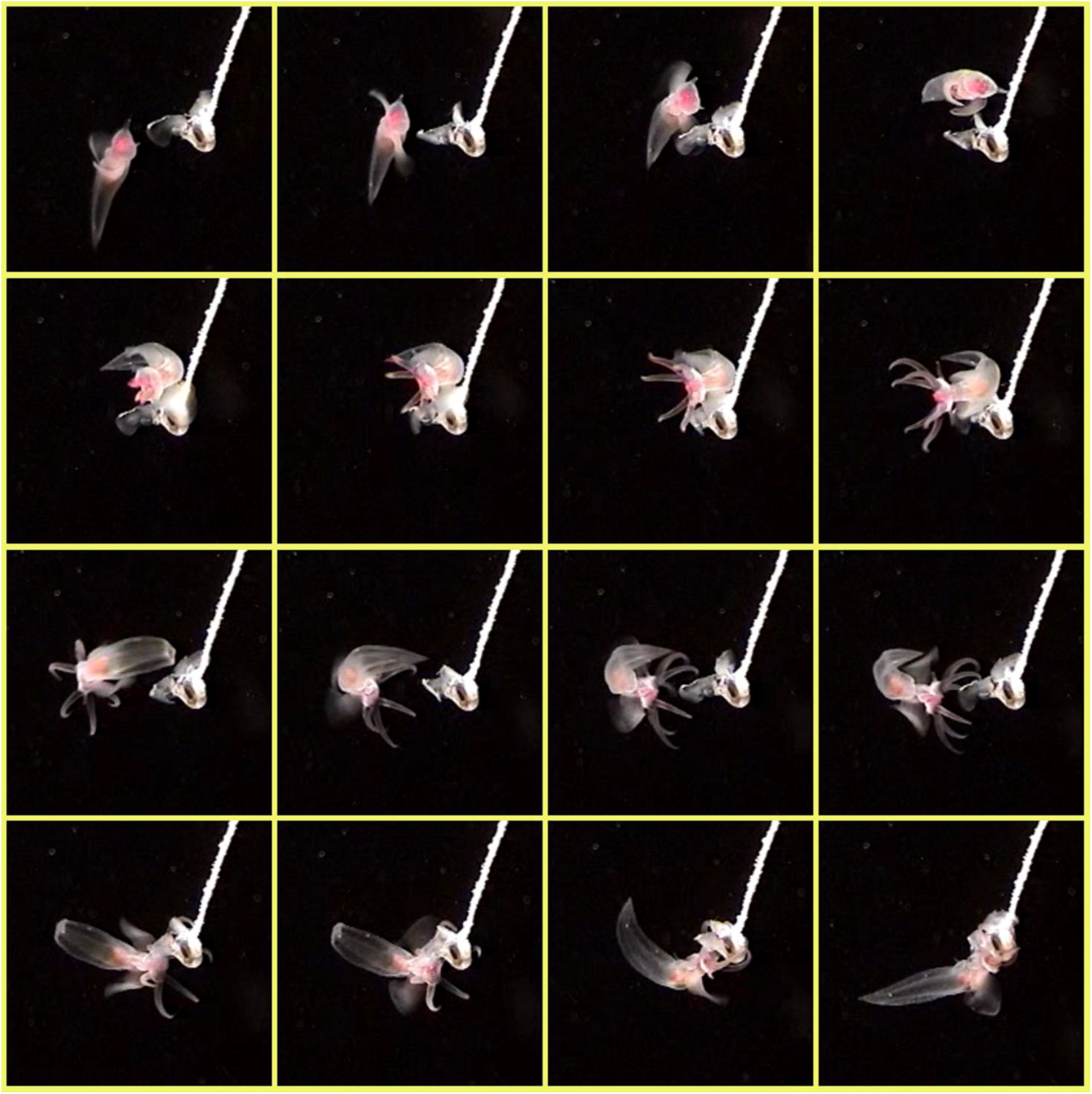
A series of sequential still images from a video depicting the feeding behavior of the sea angel *Clione limacina*. The prey, the sea devil *Limacina helicina*, is attached by its shell to a stationary stick in the center of each image (superglue was successfully used). *Clione* senses the prey and initiates the feeding behavior, protruding the prey capture tentacles and grabbing *Limacina*.

### 3.1 TH-immunoreactive neurons in the central ganglia of *Clione limacina*

The central brain in *Clione* consists of four symmetrical pairs of individual ganglia connected via central connectives and commissures – cerebral, pedal, pleural, and intestinal ganglia. TH-immunoreactive (TH-ir) neurons were found in the buccal, cerebral, and pedal ganglia in relatively small numbers (**Fig. 3a, b**; n=18). No TH-ir neurons were detected in the pleural and intestinal/visceral ganglia. Pedal ganglia had only four small-to-medium sized TH-ir neurons each, located on their ventral side, which had the size of the cell bodies between 12 and 20 μm (**Fig. 3c**). Cerebral ganglia contained 10 pairs of TH-ir neurons – 10 symmetrical cells in each ganglion (**Fig. 3d, e**). One pair of small, about 10 μm in size, TH-ir neurons was located near the cerebral commissure and was clearly observed from both dorsal and ventral sides. They were always brightly stained by TH antibody and had a single axon running into the neuropile. On their dorsal side, cerebral ganglia contained 5 medium-sized TH-ir neurons in each ganglion, with the cell bodies between 20 and 30 μm (**Fig. 3d**). There were 4 more 12-to-20 μm TH-ir neurons on the ventral side of the cerebral ganglia (**Fig. 3e**). Thus, the total number of the TH-ir neurons in the central ganglia of *Clione* was 28 (14 pairs), - significantly less than observed in the pteropod’s sister lineage represented by *Aplysia* (∼70 neurons, see (Croll, 2001).

**Figure 3.**
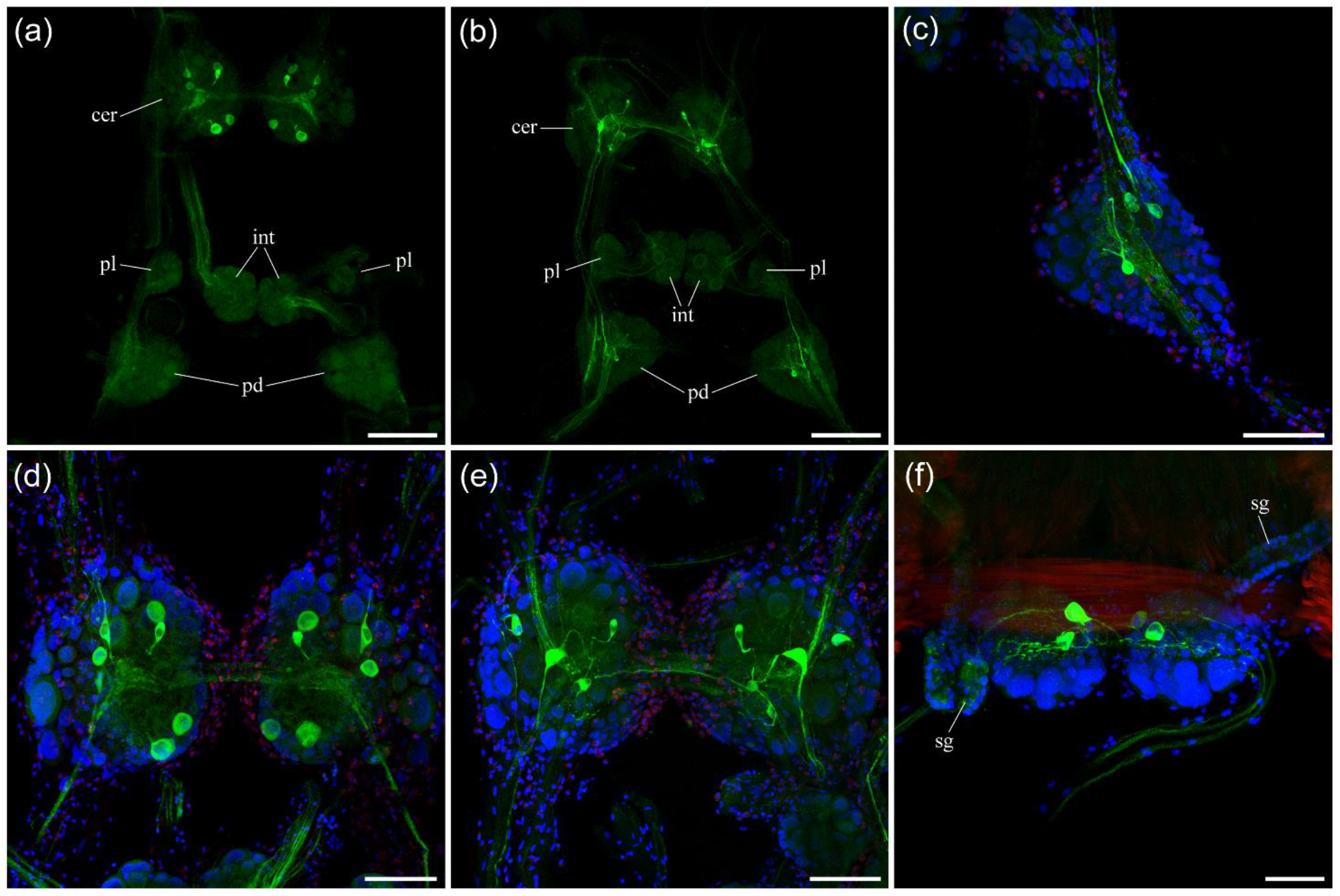
TH-immunoreactive (green) neurons in the central ganglia of *Clione limacina*. Nuclear stain by DAPI is blue. (a) Dorsal surface of the central ganglia. TH-ir neurons are seen only in the cerebral ganglia (*cer*). (b) Ventral surface of the central ganglia. TH-ir neurons in the cerebral (*cer*) and pedal (*pd*) ganglia. (c) Pedal ganglion, ventral surface, higher magnification. There are only 4 medium-sized TH-ir neurons. (d) Cerebral ganglia, dorsal surface, higher magnification. There are 6 medium-sized TH-ir neurons in each cerebral ganglion. (e) Cerebral ganglia, ventral surface. There are 5 TH-ir neurons on the ventral surface. One pair of TH-ir neurons located closer to the cerebral commissure is seen well from both dorsal and ventral sides. (f) Buccal ganglia. There are two symmetrical neurons on the ventral side. On the dorsal side - one brightly stained neuron in the left buccal ganglion, while its symmetrical cell most of the time is barely detectable. Abbreviations: *cer* – cerebral ganglia; *pd* – pedal ganglia; *pl* – pleural ganglia; *int* – intestinal ganglia; *sg* – salivary gland. Scale bars: a, b - 200 μm; c, d, e - 100 μm; f - 50 μm.

The small buccal ganglia in *Clione*, which are attached to the buccal mass, contained two symmetrical TH-ir neurons on the ventral side (Fig. 3f). On the dorsal side, one neuron was brightly stained in one (usually left) buccal ganglion. In contrast, its symmetrical neuron was barely detectable.

### 3.2 TH-immunoreactive neurons in the central ganglia of *Melibe leonina*

The CNS of *Melibe* is more centralized, compared to pteropods and *Aplysia*, and consists of four fused ganglia - two lateral pedal ganglia and two large cerebropleural ganglia connected by a short commissure (**Fig. 4a**). **Fig. 4** summarizes the distribution of TH-ir neurons in the brain of *Melibe* (n=14).

**Figure 4.**
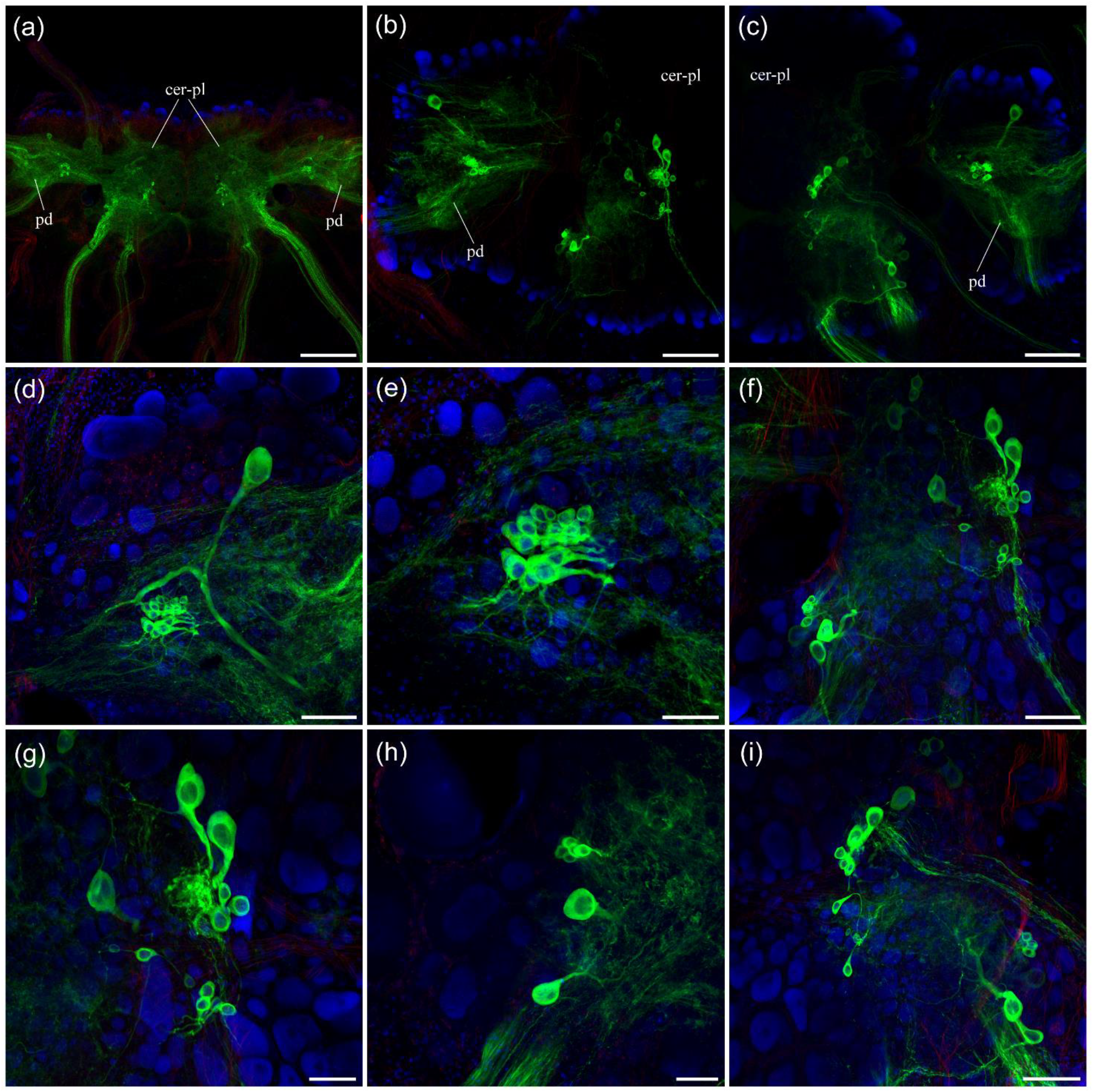
TH-immunoreactive (green) neurons in the central ganglia of *Melibe leonina*. Nuclear DAPI labeling is blue. (a) The central brain consists of fused symmetrical pedal ganglia (*pd*) and a pair of cerebropleural ganglia (*cer-pl*). (b) Left pedal and left cerebropleural ganglia; (c) right pedal and right cerebropleural ganglia. Each pedal ganglion contains one large TH-ir neuron and a cluster of small TH-ir neurons next to it. Each cerebropleural ganglion contains two separate but loose groups of TH-ir neurons. (d) Higher magnification of the right pedal ganglion with a single large TH-ir neuron and a tight cluster of small TH-ir neurons. (e) This tight cluster of small TH-ir neurons in the pedal ganglion contains between 15 and 20 cells. (f) The left cerebropleural ganglion contains roughly two groups of TH-ir neurons - one closer to the commissure and the second more posterior and laterally. (g) The group next to the commissure contains three medium-sized and 7-10 small TH-ir neurons in the left cerebropleural ganglion. (h) Higher magnification of the second group of TH-ir neurons in the left cerebropleural ganglion, which consists of two medium-sized TH-ir neurons and a cluster of 5 small TH-ir neurons. (i) Right cerebropleural ganglion with two groups of TH-ir neurons. Abbreviations: *cer-pl* – cerebropleural ganglia; *pd* – pedal ganglia. Scale bars: a - 500 μm; b, c - 200 μm; d, f, i - 100 μm; e, g, h - 50 μm.

The cerebropleural ganglia had two groups of TH-ir neurons each (**Fig. 4f, i**). One group was closer to the commissure and contained 3 medium-sized TH-ir neurons with a cell body 30 to 40 μm and a single large axon each, and 7-10 small, 15 to 20 μm, TH-ir neurons loosely positioned in that area (**Fig. 4g**). The second group was located more laterally and posterior in each cerebropleural ganglion and consisted of two medium-sized TH-ir neurons, which had the cell body size of 30 to 40 μm and a single large axon projecting toward the neuropile, and a cluster of 5 small, about 15 μm in size, TH-ir neurons (**Fig. 4h**).

The pedal ganglia contained one large TH-ir neuron in the anterior region of each ganglion and a cluster of small neurons located in the more posterior area (**Fig. 4b, c, d**). The large TH-ir pedal neuron had a size of 50 or 60 μm and a single giant axon, which then branched in the neuropile (**Fig. 4d**). The tight cluster of small TH-ir neurons contained 15 to 20 neurons, which had the size between 10 and 20 μm and a single short axon each (**Fig. 4d, e**). Thus, the number of neurons in the pedal ganglia of *Melibe* was significantly greater than that found in both *Clione* and *Aplysia*.

Even though *Melibe* does not have a buccal mass, they still retain a pair of buccal ganglia (Trimarchi and Watson, 1992; Lee and Watson, 2022). Two small buccal ganglia are positioned next to the esophagus wall on the other side from the central ganglia and near the pedal commissure that encircles the esophagus (**Fig. 5a**). There were two symmetrical small, about 10 μm in diameter, TH-ir neurons in each buccal ganglion (**Fig. 5a-c**; n=4). In addition, there was one larger, about 20 μm, TH-ir neuron consistently seen in the left buccal ganglion but not in the right ganglion (**Fig. 5b**). The total number of the TH-ir neurons in the CNS of *Melibe* was between 66 and 84 (33-42 pairs); it was greater than in *Clione* but comparable to *Aplysia*.

**Figure 5.**
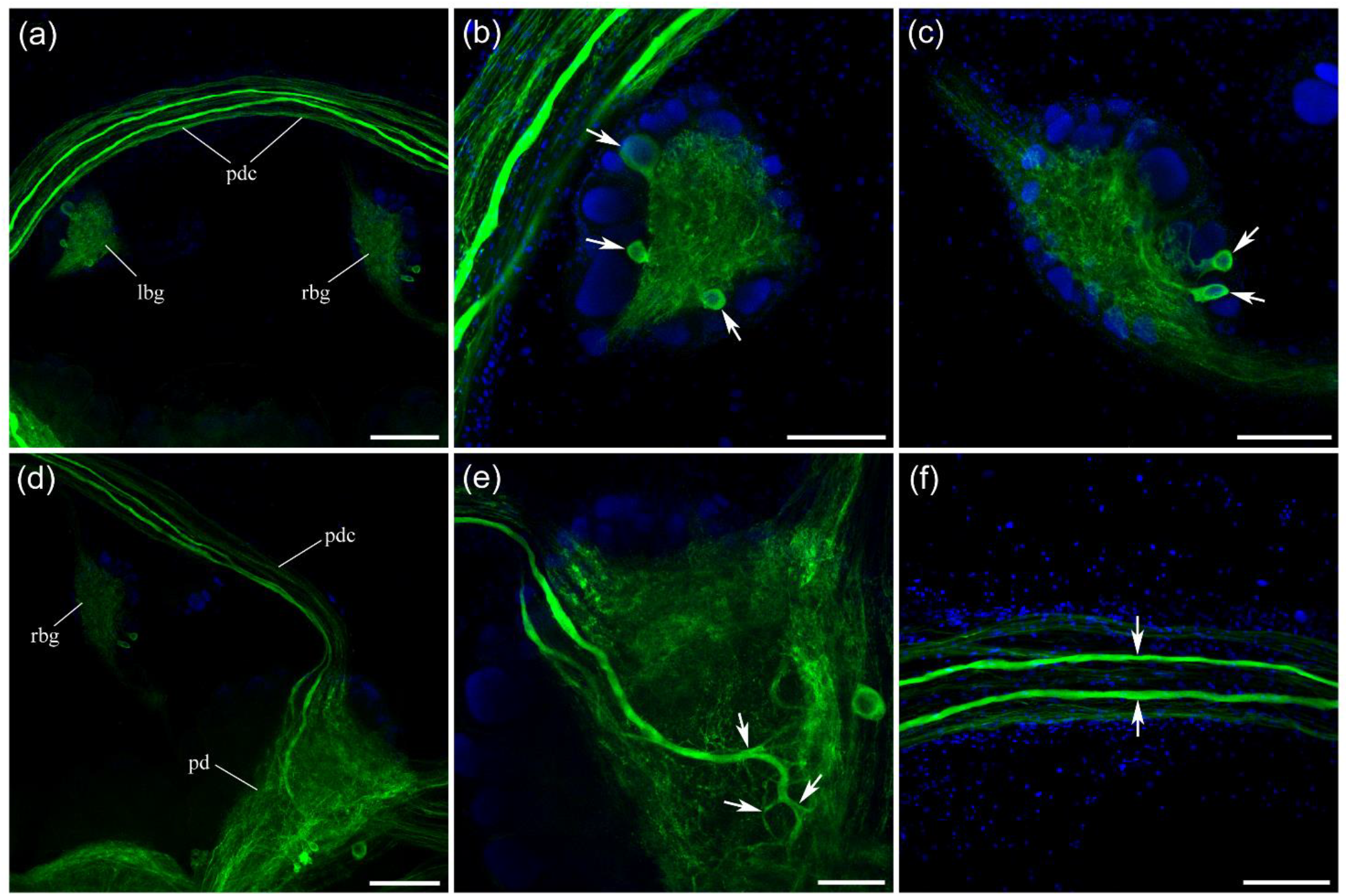
TH-immunoreactivity (green) in the buccal ganglia and pedal commissure of *Melibe leonina*. DAPI is blue (nuclei). (a) The buccal ganglia and pedal commissure are located on the opposite (from the central brain) side of the esophagus. (b) There are three TH-ir neurons (arrows) in the left buccal ganglion. (c) The right buccal ganglion contains two TH-ir neurons (arrows). (d) Pedal commissure connecting to the right pedal ganglion. (e) TH-ir axons from the pedal commissure extensively branch in the neuropile of the pedal ganglion (arrows). (f) There are two large TH-ir axons in the pedal commissure (arrows). Abbreviations: *rbg* - right buccal ganglion; *lbg* - left buccal ganglion; *pdc* - pedal commissure; *pd* - pedal ganglion. Scale bars: a, d - 200 μm; b, c, e, f - 100 μm.

### 3.3 Sensory neurons in the wings and anterior horns of the pteropod mollusc *Clione limacina*

TH-immunoreactivity revealed numerous, presumably, sensory neurons in the wings and anterior sensory tentacles (“horns”) of *Clione*. Most sensory cells in the wings were located at their distal end - at the anterior and side edges (**Fig. 6a, b**; n=18). Not many cells were seen closer to the center of the frontal edge, and there was nothing on the posterior side of the wings. There was a tight single row of numerous TH-ir sensory neurons brightly stained at the wing distal edge close to the surface (**Fig. 6a**). These TH-ir sensory neurons at the wing edge had the size of the cell body between 10 and 15 μm and sent a short single projection to the surface (**Fig. 6b**). Similar TH-ir sensory neurons were also located in the anterior sensory horns (**Fig. 6C**; n=12). They have higher concentration at the tip of the sensory tentacles and lower at their base. Each of them had the same short single projection to the surface of the tissue (**Fig. 6c**).

**Figure 6.**
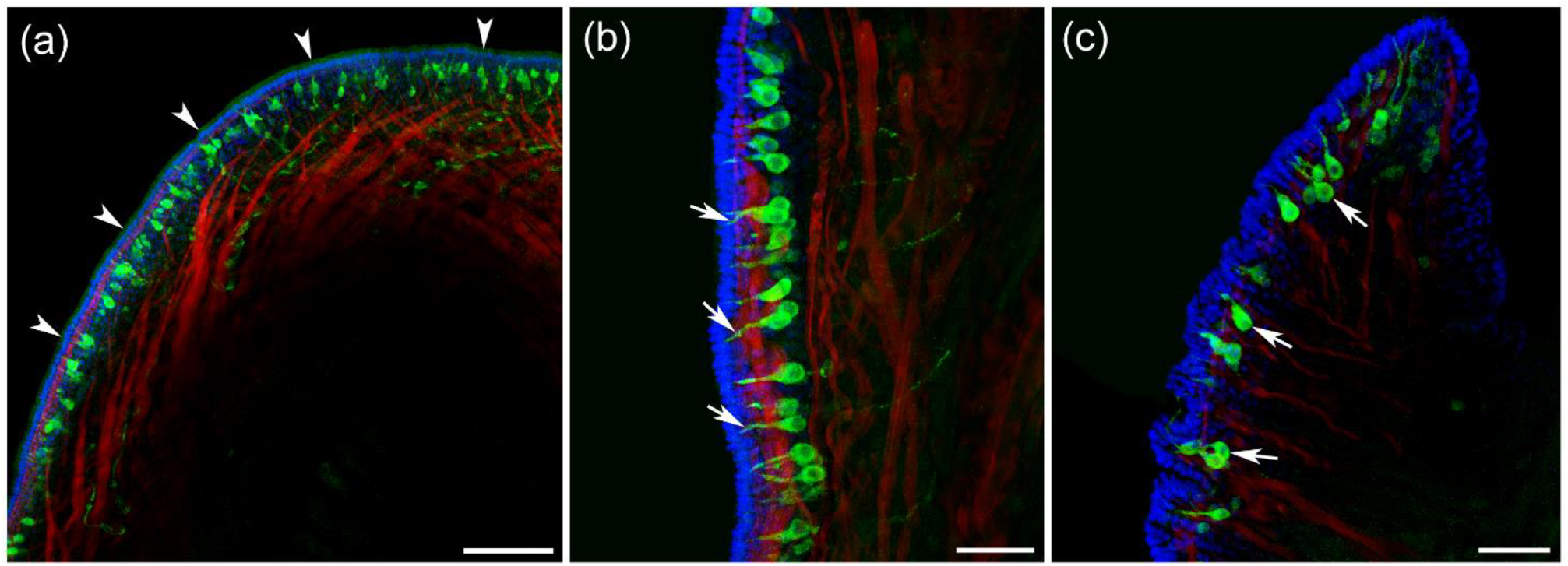
TH-immunoreactive (green) sensory neurons in the wings and anterior sensory horns of *Clione*. DAPI is blue (nuclei), while phalloidin labeling is red. The observed TH-ir neurons belong to one morphological type with the apical processes penetrating the epithelial layer. (a) The edge of the distal end of the wings (shown by arrowheads) contains a row of numerous TH-ir sensory neurons. (b) These TH-ir sensory-likw neurons at the wing edge send a short single projection each to the wing surface (indicated by arrows). (c) Similar TH-ir sensory neurons (arrows) are also located in the anterior sensory horns. Scale bars: a - 100 μm; b, c - 50 μm.

Based on our previous experience, tubulin-immunoreactivity (tubulin-IR) reveals thin neural processes much better than many other antibodies. Therefore, we used it to look at the same TH-ir sensory neurons in the periphery to obtain more details about their delicate anatomical structures. The sensory-like neurons at the edge of the wings were identified with tubulin-IR (**Fig. 7a, b**; n=14). Importantly, tubulin-IR showed that each sensory-like neuron had a single long neural process, which exited its base and then formed thicker peripheral nerves in the wings (**Fig. 7a, b**). Higher magnification also showed that each surface projection expanded at the very end, looking like a brush (**Fig. 7c**).

**Figure 7.**
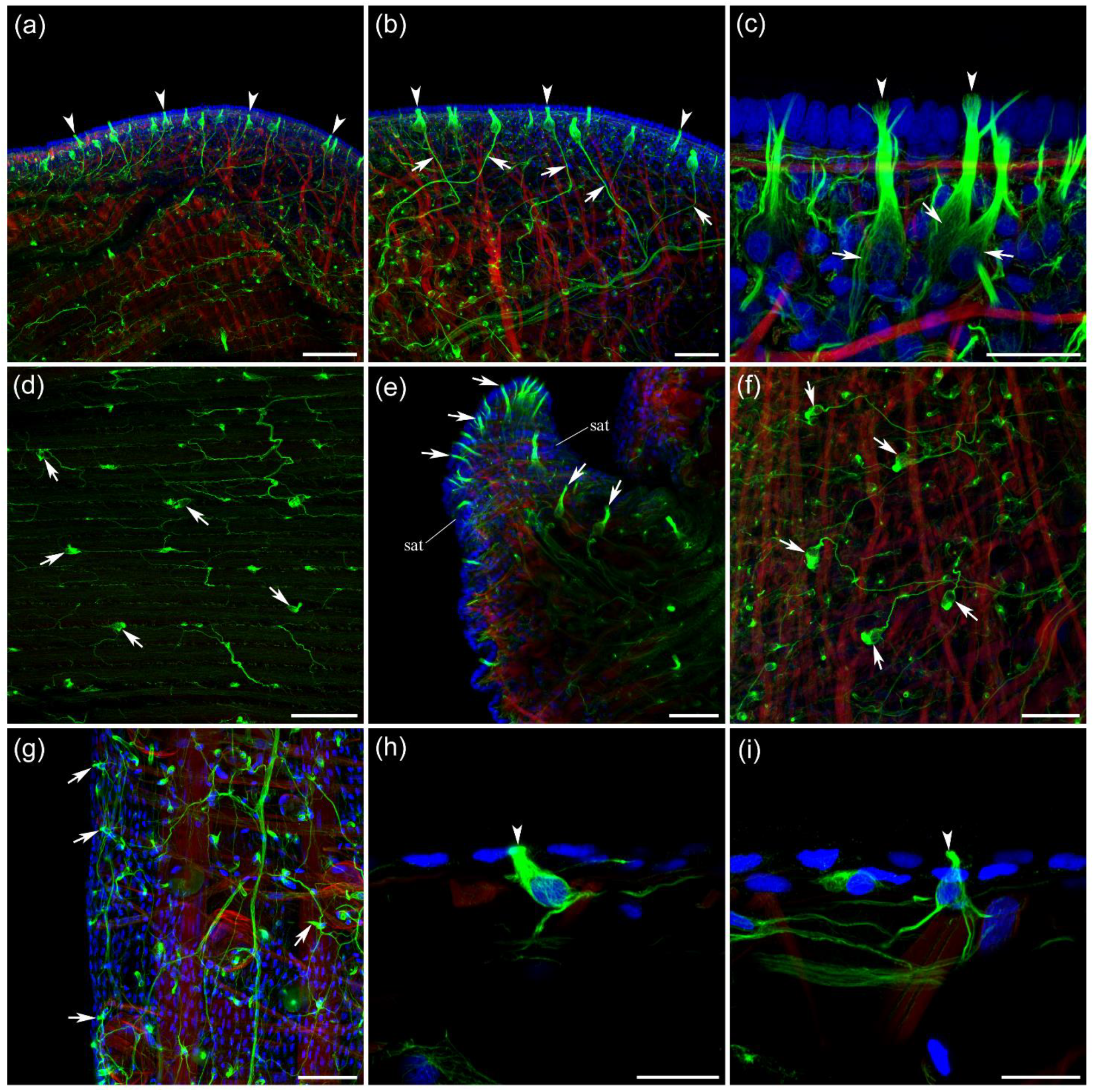
Tubulin-immunoreactivity (green) reveals the same sensory-type neural elements (as in Fig. 6) in *Clione*, but with greater details of their morphology and branching patterns. DAPI is blue (nuclei), while phalloidin labeling is red. (a) The edge of the distal end of the wing (indicated by arrowheads) contains a row of sensory neurons revealed by tubulin-IR. (b) These sensory-like neurons have a single short projection towards the surface of the wing (arrowheads show some of them) and a long neural process (arrows) each that forms thicker peripheral nerves in the wings. (c) Higher magnification shows that at the very end, near the surface, each projection expands and looks like a brush (arrowheads). Arrows indicate the cell bodies of those sensory neurons. (d) A meshwork of neurons (arrows) is also revealed on the flat surfaces of the wings, although at lower densities. These neurons might represent different subclass of cells distinct from TH-ir cells. (e) Sensory anterior tentacles (*sat*) or horns contain sensory-like neurons with a short neural projection towards the surface, at a relatively higher concentration closer to the tips of the horns. These cells are presumably the same as TH-ir neurons in Fig. 6c. (f) Similar sensory cells (arrows) and neural fibers are seen in the head skin. (g) The body wall also contains sensory-type cells (some are shown by arrows) connected by neural processes to the thicker nerves. (h, i) Many of these sensory-type neurons in the body wall, however, have multiple processes at their base. Arrowheads indicate short and thick surface projections. Scale bars: a, d, g - 100 μm; b, e, f - 50 μm; c, h, i - 20 μm.

The same sensory-like neurons were also revealed by tubulin-IR in the sensory anterior tentacles or horns - with a short neural projection toward the surface and at a relatively higher concentration closer to the tips of the horns (**Fig. 7e**; n=10). Similar sensory cells and their neural processes were observed in the skin of the head (**Fig. 7f**; n=8). The body wall also contained sensory-type cells with surface projection, which were connected by neural processes to the thicker nerves (**Fig. 7g, h, i**). Most peripheral neurons in the body wall, however, had multiple processes at their base. Many neural elements labeled by tubulin-immunoreactivity were not TH-ir, which served as a reasonable control for anti-TH antibodies. For example, effector nerves in the buccal mass of *Clione* – such as the extensively branching hook nerves that exit the buccal ganglia, innervate the hook sacs, and cause the powerful protraction of the hooks - were not TH-ir (**Fig. 8**).

**Figure 8.**
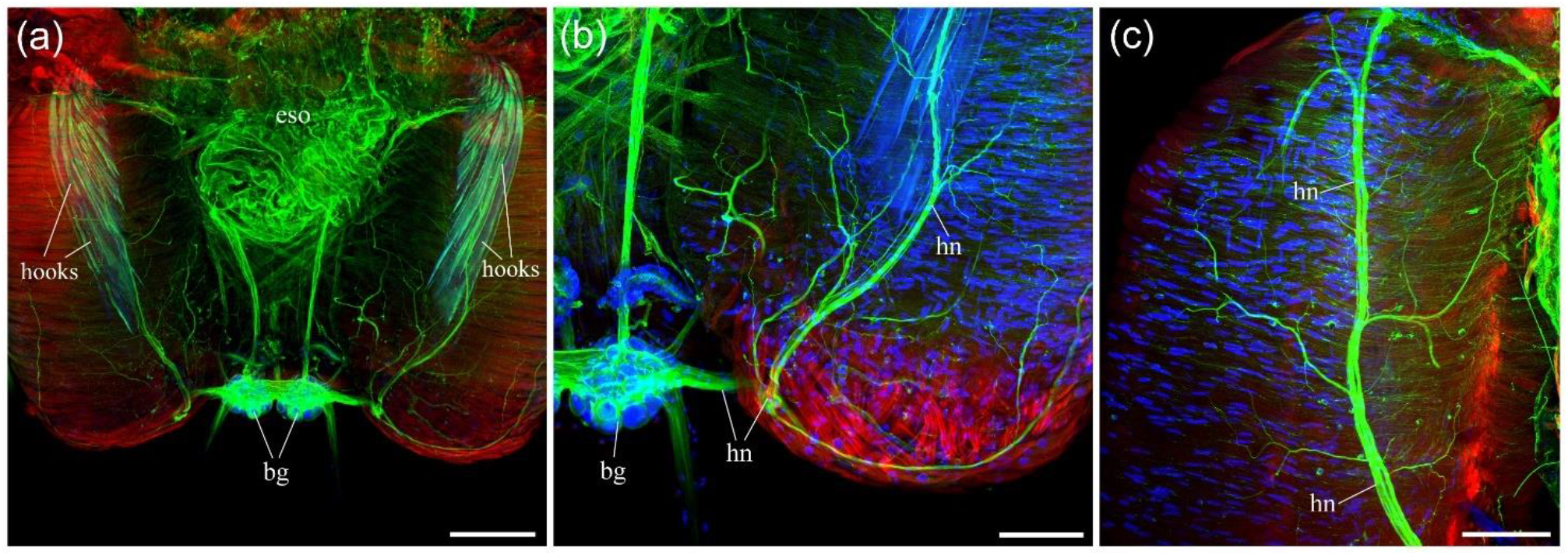
Tubulin-immunoreactivity (green) in the buccal mass of *Clione*, which does not co-localize with any TH-immunoreactivity. DAPI is blue (nuclei), while phalloidin labeling is red. (a) Isolated buccal mass, which includes a pair of muscular hook sacs and a part of the esophagus (*eso*) next to the radula, with attached buccal ganglia (*bg*). (b) Extensively branching hook nerve (*hn*) exits the buccal ganglia and innervates the base of the hook sac. (c) The hook nerve (*hn*) runs all the way to the top of the hook sac. Tubulin-IR labels buccal peripheral nerves nicely, however, these large effector nerves are not TH-ir. Abbreviations: *bg* – buccal ganglia; *eso* – esophagus; *hn* – hook nerve. Scale bars: a - 200 μm; b, c - 100 μm.

### 3.4 Sensory neurons in the wings and reduced foot of the pteropod mollusc *Limacina helicina*

Among pteropods, wings in thecosomes (like *Limacina*) are wider and more prominent in relation to their body size compared to those of Gymnosomata (like *Clione*, **Fig. 1a,b**). These differences reflect lifestyles because gymnosomes are active predators able of rapid maneuvers to hunt and capture the prey, while thecosomes with their heavy shells simply support themselves in the water column without significant directional movements (Lalli and Gilmer, 1989). *Limacina* wings contained a well-defined row of the TH-ir sensory neurons around their edges (**Fig. 9a, b**; n=4). Also, similarly to *Clione*, each sensory-like neuron had a small cell somata with a single short projection through the epithelium. However, the *Limacina* TH-ir neurons appeared to be more evenly spread around the edge of the entire wing and had lower density compared to *Clione*.

**Figure 9.**
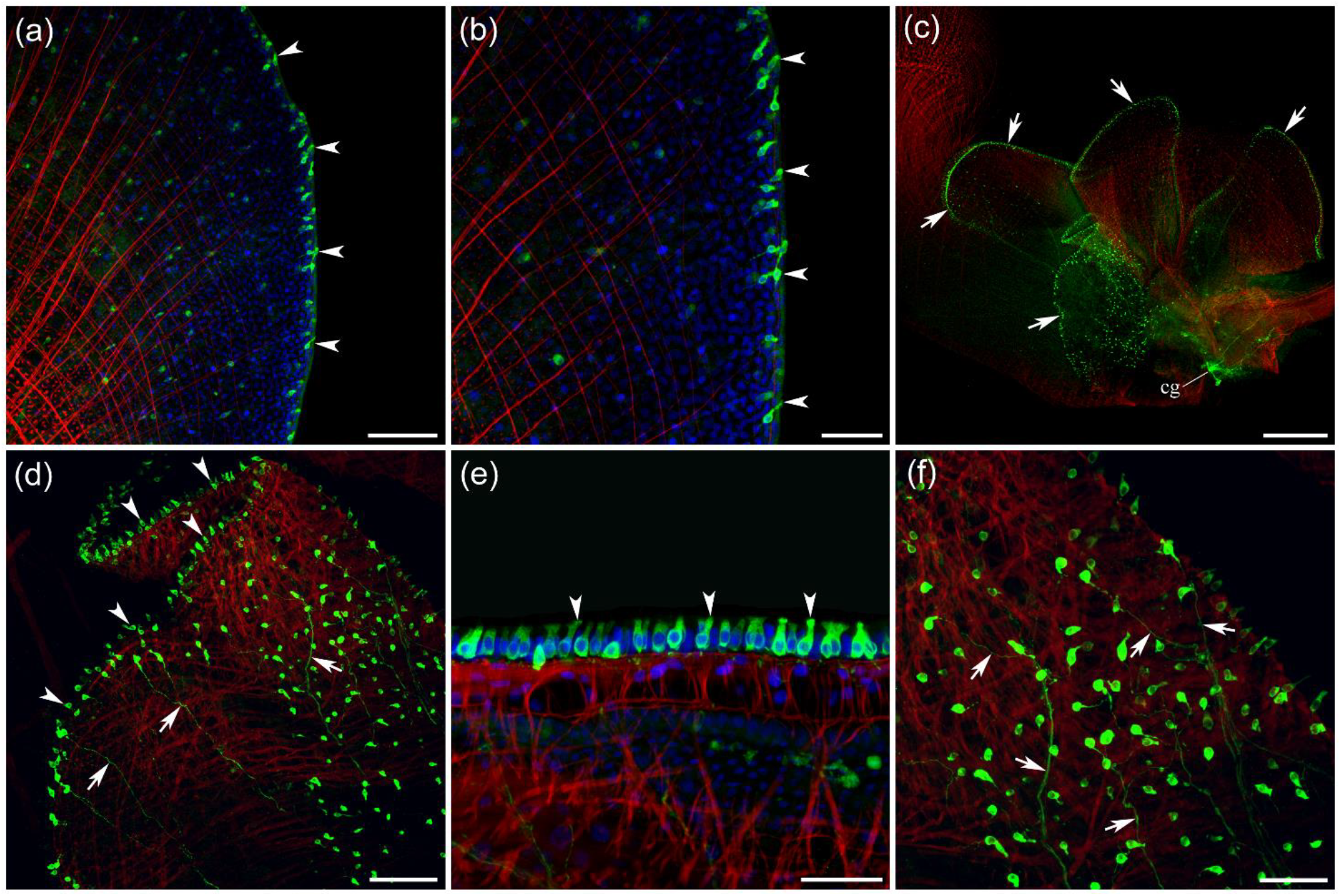
TH-immunoreactive (green) sensory neurons in the wings and foot areas of *Limacina helicina*. DAPI is blue (nuclei), while phalloidin labeling is red. (a, b) Sensory-like cells at the edge of the *Limacina* wings (indicated by arrowheads). Note the radial and circular muscle fibers in the wings labeled by phalloidin (red). (c) *Limacina* foot consists of the median lobe and lateral lobes, which look like tissue folds between wings. Those pedal folds contain at their very edge rows of TH-ir cells (arrows). (d) TH-ir sensory neurons in the fold of the median lobe – arrowheads point to the edge of the fold, and arrows indicate the long TH-ir processes. (e) Tight row of TH-ir sensory cells at the edge of the lateral lobe of the foot. Arrowheads point to the short sensory projection from the cell body to the tissue surface. (f) TH-ir sensory-like cells of the foot tissue, arrows point to the multiple neural processes that connect sensory-like cells with the underlying neural network. Abbreviations: *cg* – central ganglia. Scale bars: a, d - 100 μm; b, f - 50 μm; c - 500 μm; e - 30 μm.

The reduced foot in *Limacina* consists of a median lobe and two narrower lateral lobes, which are all located between two large wings, and are involved in filtering the food and brinning it to the mouth. Each foot lobes contained at their outer edge a dense row of TH-ir cells (**Fig. 9d, e**) with the apical sensory-like region (**Fig. 9c-f**; n=4). Long TH-ir neural processes connected these sensory cells with the underlying neural network (**Fig. 9d, f**). Each TH-ir sensory neuron had a short sensory projection from the cell body to the tissue surface, which is typical for this cell type.

In the wings of *Limacina*, tubulin-immunoreactivity (similar to *Clione*) revealed a significantly greater number of multipolar neurons, which were not TH-immunoreactive (**Fig. 10**; n=4), suggesting the greater complexity of neural networks in the wings (and also served as a proper control for the specificity of anti-TH antibodies).

**Figure 10.**
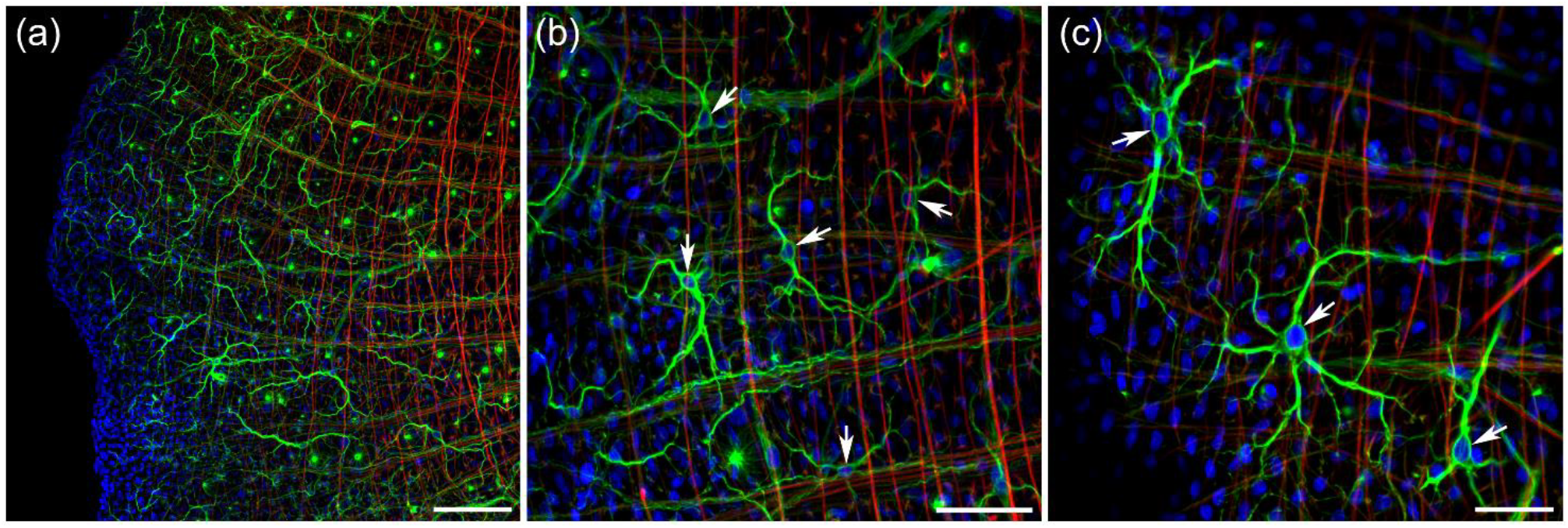
Tubulin-immunoreactivity (green) in the wings of *Limacina* shows more neural elements than TH-immunoreactivity. DAPI is blue (nuclei), while phalloidin labeling is red. (a) The surface of the wing with numerous neural elements. (b, c) Tubulin-IR labels multipolar neurons (shown by arrows), which are not TH-ir. Scale bars: a - 100 μm; b - 50 μm; c - 30 μm.

### 3.5 TH-immunoreactive sensory elements in the rhinophores of *Melibe leonina*

*Melibe* has two symmetrical ear-like chemosensory structures on the oral hood of its head called rhinophores. TH-immunoreactivity was abundant in *Melibe* rhinophores, including actively branching TH-ir nerves, which covered the entire sensory organ (**Fig. 11a**; n=12). The thickest branch of TH-ir nerves ran across each rhinophore and approached the rhinophore groove on the outer side (**Fig. 11b**). Many TH-ir sensory neurons were located at the edge of the rhinophore groove (**Fig. 11c, d**). The density of TH-ir sensory cells was much higher at the edge of the groove than in the rest of the rhinophore (**Fig. 11e, f**). These TH-ir neurons were connected to the nerves by thin neural processes (**Fig. 11c-e, g-i**) and were organized in clusters (**Fig. 11d-f, i**). Each TH-ir sensory neuron had a very thin and relatively short projection from the cell body to the tissue surface (**Fig. 11f, i**). This projection was typical for this subtype of TH-ir sensory cells in all studied species. However, in *Melibe*, these projections were much thinner and more prolonged than in *Clione* and *Limacina* (both of these species had much thicker and shorter projections).

**Figure 11.**
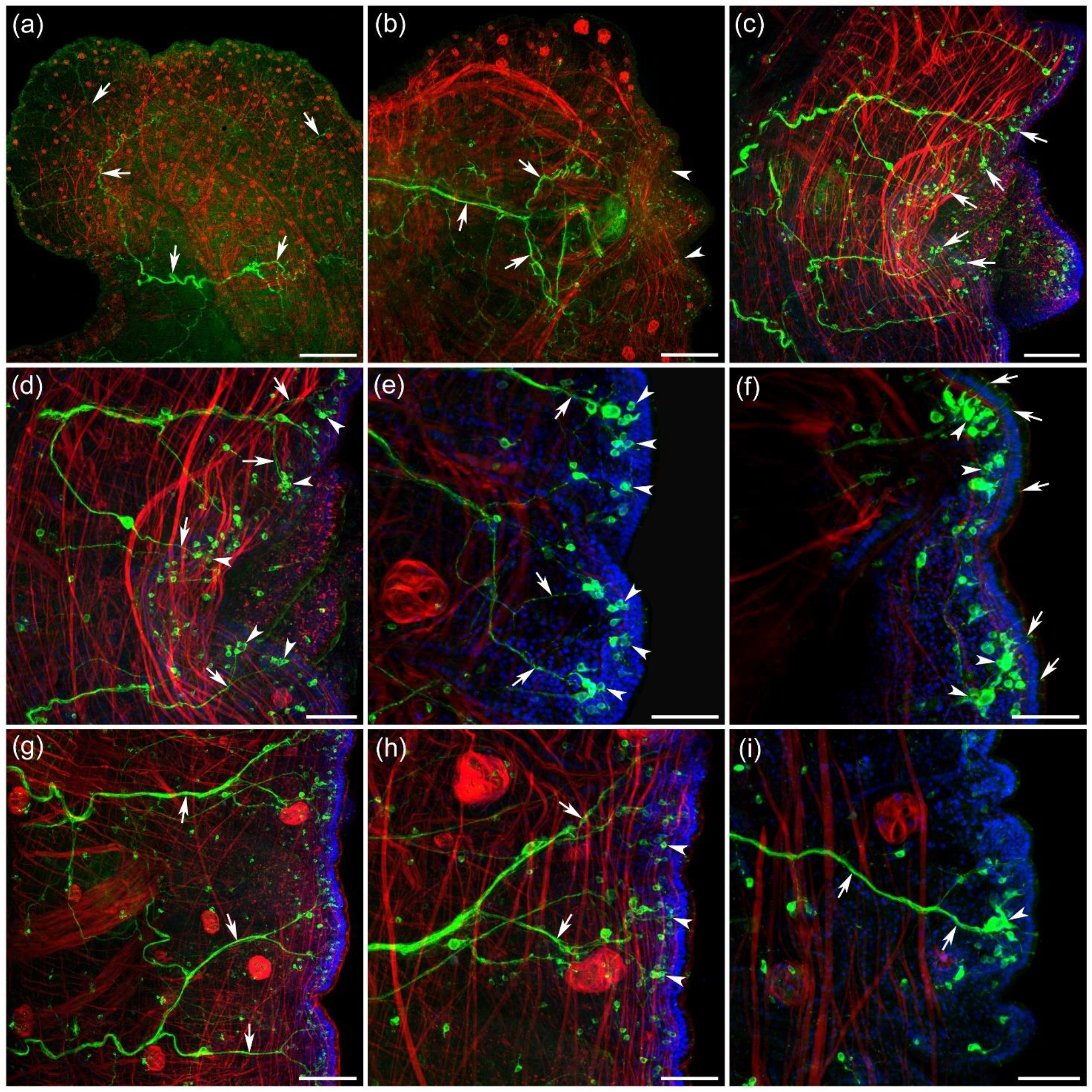
TH-immunoreactive (green) sensory neurons in the rhinophores of *Melibe leonina*. DAPI is blue (nuclei), while phalloidin labeling is red. (a) *Melibe* rhinophores have branching TH-ir nerves (arrows) covering the entire sensory organ. (b) The thickest TH-ir nerve (arrows) approaches the rhinophore groove (indicated by arrowheads). (c) Many sensory-like TH-ir neurons (arrows) are located at the edge of the rhinophore groove. (d) Those TH-ir neurons are connected to the nerve by thin neural processes (arrows). Note that sensory neurons are grouped in clusters (arrowheads). (e) The densoty of TH-ir sensory cells (arrowheads) is higher at the edge of the groove than over the rest of the rhinophore. Arrows point to the thin TH-ir processes connecting sensory neurons to the nerves. (f) Note thin projections (arrows) from the neuronal somata, through the epithelium to the surface. These neurons tend to cluster in groups (arrowheads). (g) The TH-ir nerve branches (arrows) innervate the entire area of the rhinophores, not only its groove area. (h) The density of the TH-ir sensory neurons (arrowheads) at the edge of the rhinophores is lower outside of the rhinophore groove. Arrows point to the branching nerves, which are formed by thin neural processes originating from the sensory neurons. (i) A cluster of sensory neurons (arrowhead) is connected to the nerve branch (arrows). Note also the thin, short projections from the TH-ir sensory neurons to the surface of the rhinophore. Scale bars: a - 500 μm; b - 200 μm; c, g - 100 μm; d, e, f, h, i - 50 μm.

In the *Melibe* rhinophores, tubulin-IR revealed the same type of neural elements as TH-immunoreactivity with broadly branching nerves (**Fig. 12a**). Upon reaching the edge of a rhinophore, tubulin-ir terminals innervated sensory-like clusters of cells (**Fig. 12b, c**). In *Melibe* most peripheral neural elements labeled by tubulin-ir were also TH-immunoreactive.

**Figure 12.**
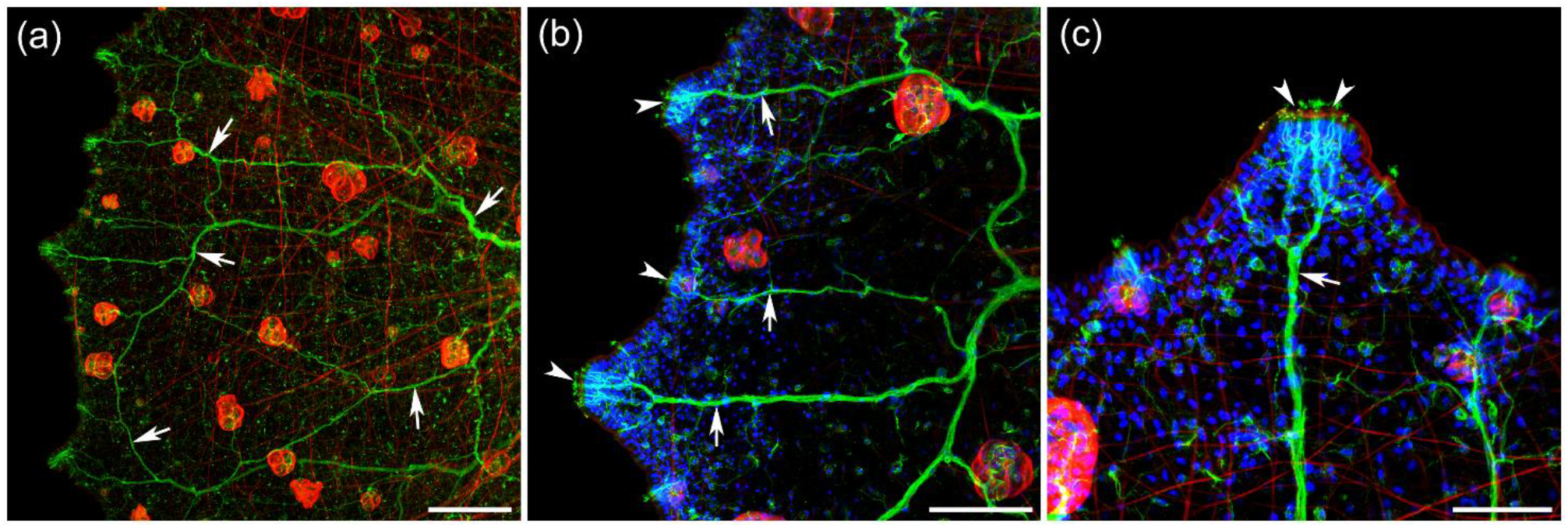
Tubulin-immunoreactivity (green) reveals the same sensory neural elements in the *Melibe* rhinophores as labeled by TH-ir (Fig. 11). DAPI is blue (nuclei), while phalloidin labeling is red. (a) Actively branching nerves (arrows) in the rhinophores labeled by tubulin-ir. (b) Upon reaching the edge of a rhinophore tubulin-IR terminals (arrows) innervate the sensory clusters (arrowheads). (c) Higher magnification of the sensory cluster (arrowheads) at the edge of a rhinophore that is innervated by the tubulin-ir nerve branch (arrow). Scale bars: a - 200 μm; b - 100 μm; c - 50 μm.

### 3.6 TH-immunoreactive sensory elements in the oral hood of *Melibe leonina*

*Melibe* has a unique large expandable oral hood used to capture small planktonic animals, with two rows of cirri or protrusions (papillae) around its outside edge. Those protrusions are presumably sensory structures used to detect prey. One row carries much longer protrusions, and one row has shorter ones (**Fig. 1c**). TH-immunoreactivity was abundant in these sensory structures, most notably labeling their thick nerves (**Fig. 13a-c**; n=14). Longer papillae had one single TH-ir non-branching nerve in the center, running to their thin tips (**Fig. 13a**). The shorter papillae had intensively branching TH-ir nerve inside (**Fig. 13b, c**). Hundreds of small TH-ir neurons were found on the surface of both types of these structures (**Fig. 13d-f**).

**Figure 13.**
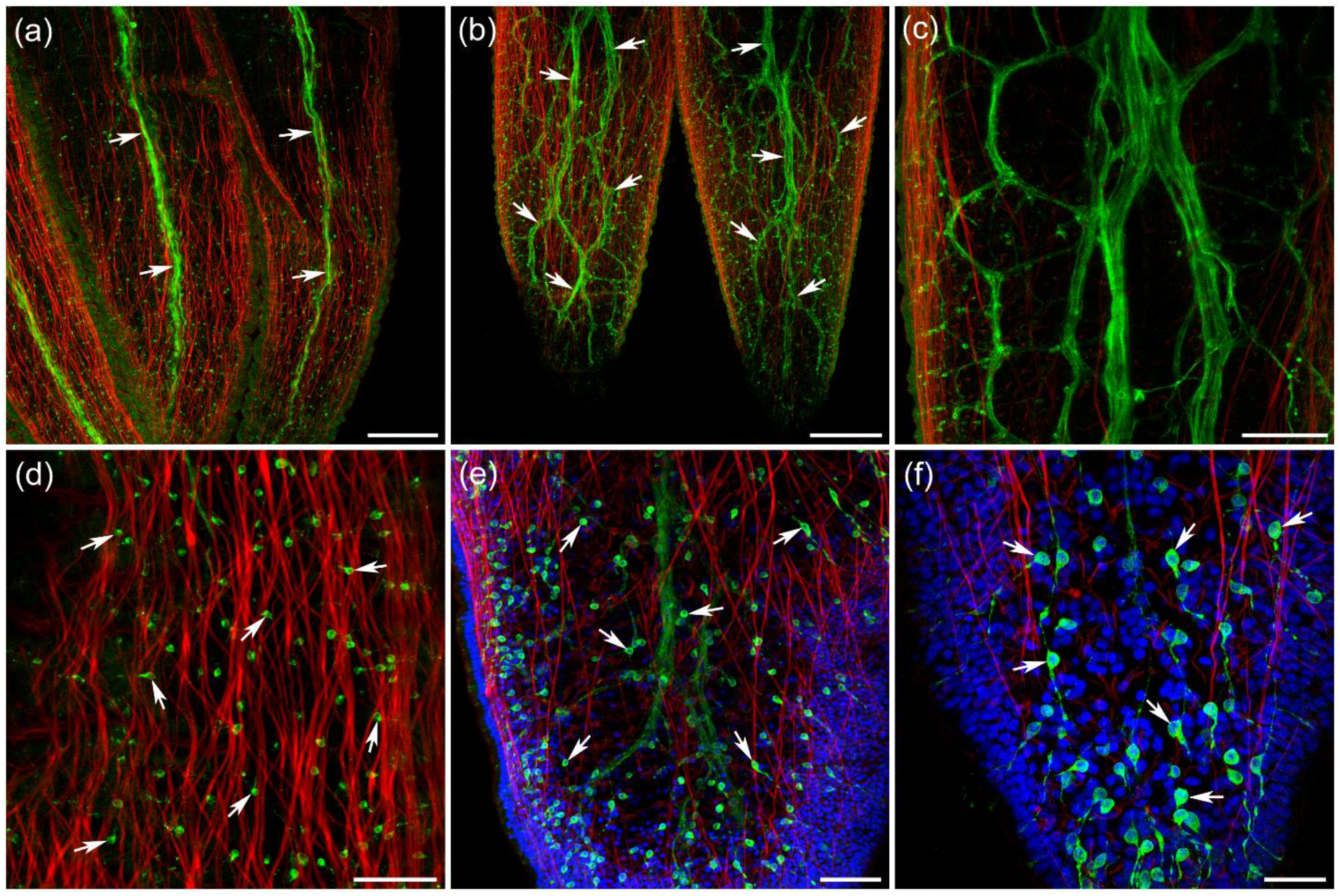
TH-immunoreactivity (green) in the oral hood sensory cirri of *Melibe leonina*. DAPI is blue (nuclei), while phalloidin labeling is red. (a) There are two rows of sensory protrusions along the oral hood edge – one row of longer and one row of shorter protrusions. Longer protrusions have one single TH-ir non-branching nerve (arrows) in the center of each structure. (b) The shorter protrusions have actively branching TH-ir nerve (arrows). (c) The branching TH-ir nerve inside shorter protrusions at higher magnification. (d) TH-ir neurons (arrows) on the surface of the longer protrusions. (e) TH-ir neurons (arrows) on the surface of the shorter protrusions. (f) High magnification of TH-ir neurons (arrows) on the surface of a shorter protrusions. Note thin neural processes exiting each neuron (mostly bipolar). Scale bars: a, b - 200 μm; c - 100 μm; d, e - 50 μm; f - 25 μm.

Tubulin-IR labeled the same nerves in the oral hood cirri as TH-immunoreactivity - a single non-branching nerve inside each more extended protrusions and the actively branching nerve in the shorter extensions (**Fig. 14a-c**). All nerves labeled by tubulin-IR in the protrusions also contained TH-ir processes. Tubulin-IR revealed more clearly (than TH-ir) numerous small peripheral sensory projections from the nerves to the surface. Thin neuronal processes connected the central non-branching nerves of the longer protrusions with numerous sensory clusters on the surface (**Fig. 14d, e**). Tubulin-IR unmistakably identified those sensory clusters (with greater microanatomical details) on the surface of both types of cirri.

**Figure 14.**
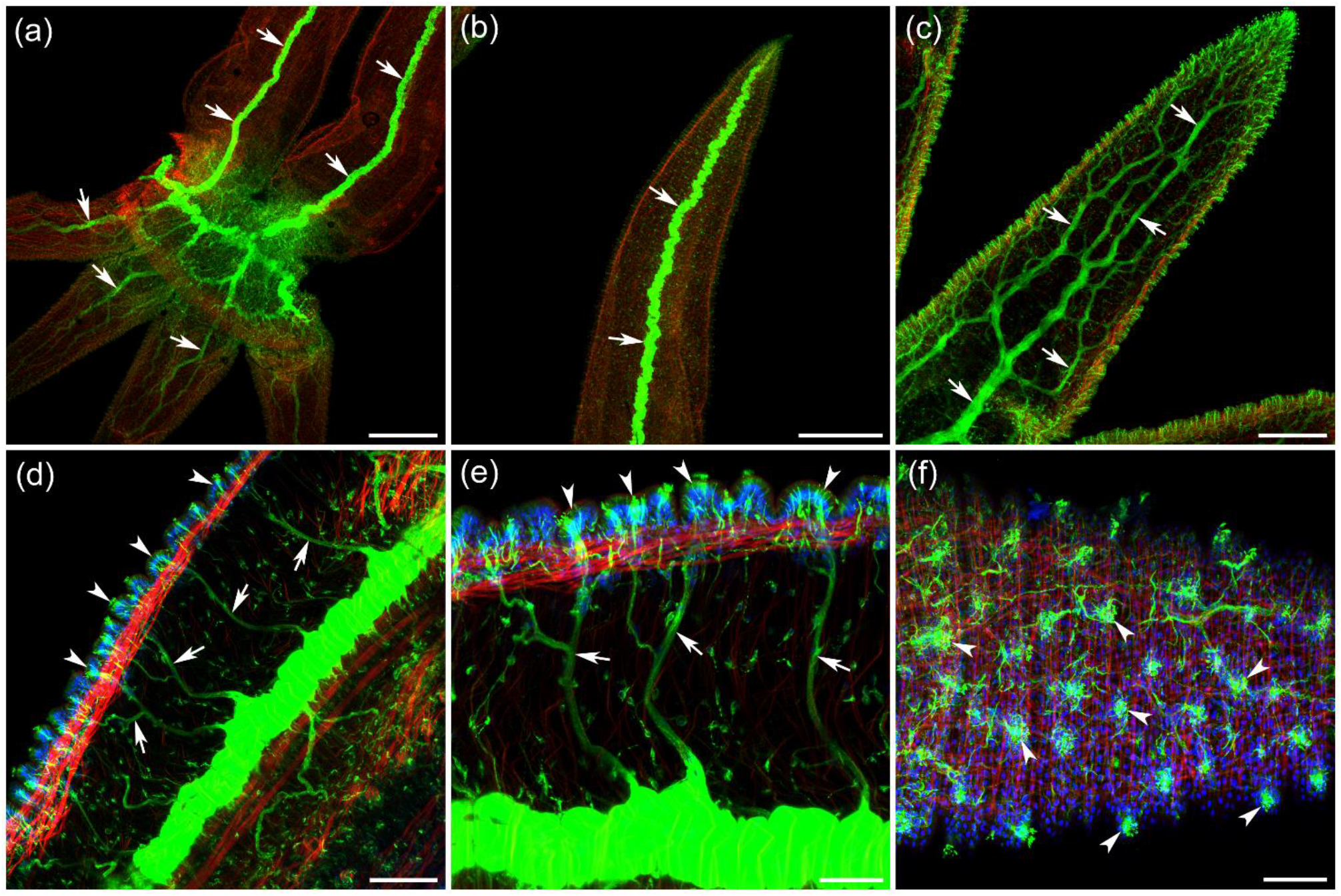
Tubulin-immunoreactivity (green) reveals the same neural elements in the *Melibe* oral hood sensory cirri. DAPI is blue (nuclei), while phalloidin labeling is red. (a) There are two rows of protrusions around the edge of the hood, which all are innervated by tubulin-ir nerves (arrows). (b) One row consists of longer extensions with a single non-branching nerve inside each protrusion (arrows). (c) The second row consists of shorter extensions with the actively branching nerve (arrows). (d) Thin neural fibers (arrows) exit the single non-branching nerve of the longer cirri and reach the surface where they innervate the sensory clusters (arrowheads). (e) Higher magnification of the neural fibers (arrows) that connect the central thick nerve with the sensory clusters (arrowheads) on the surface. (f) Sensory clusters on the surface of a shorter protrusions. Scale bars: a, b - 500 μm; c - 200 μm; d - 100 μm; e, f - 50 μm.

## 4 DISCUSSION

With over 30,000 species, euthyneuran gastropods represent one of the most diverse lineages in Mollusca, and play significant ecological roles in aquatic and terrestrial environments (Ferrari and Hautmann, 2022). The presence of large, identified neurons in representatives of Euthyneura justified the early focus on dopamine signaling with TH-ir or glyoxylate-induced histofluorescence mapping. Although there is no 1:1 correspondence between both techniques (assuming their apparently different sensitivity), there is a reasonable consensus that TH-ir is more sensitive than histofluorescence, and predominantly labels DA-containing cells both in sea slugs (Croll, 2001) and pulmonate molluscs (Elekes et al., 1991; Elekes, 1992; Vallejo et al., 2014). Although other catecholamines are possibly present in gastropods, their concentrations are much lower than DA, and only a small fraction of neurons (a few individual cells at maximum) might contain other types of catecholamines (Miller, 2020). Of note, earlier biochemical tests suggesting the presence of noradrenaline or adrenaline (Osborne and Cottrell, 1971; Osborne et al., 1975; McCaman et al., 1979; Kiehn et al., 2001) need to be cross-validated with alternative, modern, more sensitive bioanalytical techniques. Thus, we refer to TH-ir cells in this discussion as dopaminergic systems.

### Central dopaminergic neurons across Euthyneura: Overview

Extensive comparative mapping in freshwater pulmonate molluscs (superorder Hygrophila) led to the discovery of an asymmetric giant dopaminergic neuron in the right pedal ganglion of *Lymnaea* (Winlow et al., 1981) and other Lymnaeidea with the dextran shell coiling. In snails with the sinistral shell coiling (e.g., *Helisoma, Biomphalaria, Planorbis*, etc.), the giant dopaminergic cell is present in the left pedal ganglion (Pentreath et al., 1974; Osborne et al., 1975; Kiehn et al., 2001; Vallejo et al., 2014). Clearly, these giant dopaminergic neurons (∼200-300 μm) are homologous cells known to be involved in the control of respiratory behaviors as parts of the respiratory central pattern generator (Moroz, 1990; Syed et al., 1990; Moroz, 1991; Syed and Winlow, 1991; Moroz and Winlow, 1992b; Winlow et al., 1992). Their distant homologs have not been identified outside Hygrophila, for example, among land pulmonates such as *Helix* or *Limax* (orders Stylommatophora and Systellommatophora, respectively); *Clione, Aplysia, Melibe*, and *Pleurobranchaea* also do not have such giant dopaminergic neurons. Cell-type-specific homologization and functions of small dopaminergic neurons in the gastropod pedal ganglia are presently unknown.

More likely, buccal and cerebral ganglia contain some subsets of homologous neurons, but their tiny sizes prevent precise homologization across species (Elekes et al., 1991; Elekes, 1992; Croll, 2001; Croll et al., 2001; Croll et al., 2003; Miller, 2020).

### Comparing dopaminergic systems between nudipleuran and pteropod molluscs

Nudipleura (represented in this discussion by two clades leading to *Melibe* and *Pleurobranchaea*, respectively) is the most basally branched lineage of Euthyneura, yet their representatives have one of the most prominent centralizations of neurons, reflecting the parallel evolution of brains as differently fused ganglia architecture (Moroz, 2009). There are similarities between the distribution of dopaminergic neurons in *Melibe* and *Pleurobranchaea. Pleurobranchaea* is larger and has more neurons in the CNS than *Melibe* (see also (Boyle et al., 1983)); and this situation is reflected by comparing numbers of TH-ir neurons.

Very small buccal ganglia in the passive plankton-eater *Melibe* contain two pairs of small symmetrical TH-ir neurons and one larger asymmetrical TH-ir neuron in the left buccal ganglion. In contrast, the opportunistic predator *Pleurobranchaea* has 7 TH-ir neurons in the buccal ganglia (Brown et al., 2018). The pedal ganglia in *Melibe* contained one large TH-ir neuron in the anterior region of each ganglion and a tight cluster of small neurons located in the posterior area – slightly different distribution than in *Pleurobranchaea*, which has two loose clusters of small TH-ir neurons on the ventral side of pedal ganglia (Brown et al., 2018). Most TH-ir neurons in both species were located in the sizeable cerebro-pleural ganglion, with about 20 symmetrical pairs in *Melibe*.

The marine pteropod *C. limacina* represents the sister lineage to the well-studied sea slug, *Aplysia californica* (Pabst and Kocot, 2018). Both species have a similar arrangement of the CNS with *separate* buccal, cerebral, pleural, pedal, and intestinal (=abdominal) ganglia connected via several connectives. The number of neurons, and dopaminergic cells in the central ganglia of *Clione* is the smallest than in all other gastropods discussed here, including *Aplysia* (Croll, 2001). The overall smaller number of neurons in the pteropod CNS can viewed as an adaptation to the pelagic lifestyle with cost-efficient energetic demands. For example, the overall number of serotonin-immunoreactive neurons in the central ganglia of *Clione* is significantly less than that found in *Aplysia*, while the general distribution of those serotonergic neurons showed remarkable similarities with several cases of homologization (Satterlie et al., 1995).

Our results on *Clione limacina* confirm and extend the previous investigation of catecholaminergic neurons that was conducted using glyoxylate-induced histofluorescence method (Kabotyanski and Sakharov, 1989; Kabotyanskii and Sakharov, 1990), with the predominant localization at the periphery, primarily in the chemosensory regions of wings (a modified foot), oral cones, and the esophagus (Kabotyanskii and Sakharov, 1990; Voronezhskaya and Kabotyanski, 1991).

The general pattern of the distribution of central dopaminergic neurons also showed some similarities between *Aplysia* and *Clione*. Yet, only 3 TH-ir neurons were found in the buccal ganglia of *Clione*, while *Aplysia* had 9 buccal neurons. Most TH-ir somata were located in the cerebral ganglia and some in the pedal ganglia, with approximately 3 times more TH-ir neurons in *Aplysia* (Croll, 2001) than in *Clione*. Both species didn’t have any TH-ir somata in the pleural and intestinal (abdominal) ganglia; no TH-ir neuron somata were present in the separate visceral ganglion of *Pleurobranchaea*.

However, in freshwater pulmonate molluscs (Hygrophila), there is a number of dopaminergic neurons in parietal/visceral ganglia (homologous structures to the abdominal ganglion in *Aplysia*) as summarized in (Elekes et al., 1991; Elekes, 1992; Voronezhskaya et al., 1999; Vallejo et al., 2014). This distribution suggests lineage-specific innovations and/or migratory events of ancestral dopaminergic neurons to these structures either from the cerebral ganglia or the periphery.

### Peripheral dopaminergic neurons in Euthyneura

All studied euthyneurans contain a relatively small number of central dopaminergic neurons - usually less than 100 cells or ∼0.5-1% of the total neuronal population in the CNS. In contrast, the number of dopaminergic neurons at the periphery is astonishing (usually >1,000-5,000), and these neurons likely belong to different subclasses, with the predominance of cells with sensory-like morphology both in pteropods and other studied Euthyneura (Faller et al., 2008; Brown et al., 2018; Miller, 2020; Norekian et al., 2024). The origin and evolution of such distinct organizations in gastropods are unknown, and more extensive comparative studies are needed in both different molluscan classes and sister spiralians. These dopaminergic cells apparently represent a significant part of the peripheral sensory systems, perhaps in addition to nitrergic neurons in grazers (e.g., *Aplysia, Lymnaea, Helisoma*, and others (Moroz and Gillette, 1995; Moroz, 2000; 2006), but not in predators such as *Clione* (Moroz et al., 2000) and *Pleurobranchaea* (Moroz and Gillette, 1996)). Most dopaminergic peripheral sensory neurons have a similar basic structure – the cell body located in the subepithelial layer with one single projection towards the surface and one long axon on the opposite side forming with similar axons, the afferent nerves running towards the central ganglia. In *Clione* and *Limacina*, these putative sensory (TH-ir) neurons are primarily located at high densities in the most strategically important areas for receiving sensory information – for example, edges of the wings. In a fast-moving *Clione*, they are predominantly on the frontal edges of the wings, which mainly contact the incoming water flow (as mechanoreceptors) or chemosensory signals from the environment. They are also found on the anterior sensory tentacles (“horns”) on the top of *Clione* head and in the esophagus (Kabotyanskii and Sakharov, 1990). In slower, non-directional swimmers, like *Limacina*, TH-ir sensory neurons are found over the entire edge of the wide wings and also on the lobes of the reduced foot between the wings where the food is filtered.

Although the *Melibe* anatomy is different than that in *Pleurobranchaea* or *Clione*, the structure of the peripheral dopaminergic sensory system is similar at its basic level. TH-ir sensory neurons in *Melibe* rhinophores look more like in the *Pleurobrachaea* veil, with a relatively long and very thin projection to the surface (Norekian et al., 2024). In addition, those TH-ir sensory-like neurons in both species are grouped in clusters, and their axons join to form afferent nerves that run to the central ganglia. The unique sensory protrusions on the edge of the feeding hood in *Melibe* also have numerous TH-ir sensory neurons near the surface, whose axons then join to form nerves projecting into the head. In *Clione* and *Limacina*, however, they are slightly different – those projections are thicker and shorter.

In summary, dopaminergic cells apparently represent a significant part of the peripheral chemosensory systems, overlapping with nitrergic neurons in grazers (Moroz, 2006) but not carnivore gastropods. Dopamine can also mediate or modulate other sensory modalities (e.g., mechanoreception), and functional studies on peripheral neural circuits are highly desirable (Carrigan et al., 2015; Horváth et al., 2020; Brown et al., 2023).

### Functional and evolutionary significance of monoaminergic systems in Euthyneura

How evolutionary conservative functions of DA systems across molluscs is unclear. The role of DA in feeding is the best-studied paradigm in *Aplysia*, with reward signaling in central circuits (Brembs et al., 2002). Specifically, a few small identified dopaminergic cells in the buccal and cerebral ganglia are well-characterized as interneurons, driving feeding motor patterns (Rosen et al., 1991; Teyke et al., 1993), often via interactions with other interneurons and by directly activating buccal motoneurons (Kabotyanski et al., 1998; Kabotyanski et al., 2000). These modulatory dopaminergic cells also contribute to the operant conditioning memory (Momohara et al., 2022) as components of the feeding enjection program and an indication that the food was uneatable (Miller, 2020). It is reasonable to suggest that homologs of these dopaminergic, feeding-related interneurons are present among those few TH-ir cells discovered in the buccal and cerebral ganglia of *Clione* and *Melibe*.

The pharmacological data confirm that the central feeding circuit received DA-mediated input from chemoreceptive cells located in the esophagus, mouth area, rhinophores, and parts of the foot and head epithelia (Martínez-Rubio et al., 2009; Brown et al., 2018). Thus, in gastropods studied so far, most dopaminergic neurons are located in the peripheral nervous systems and associated with primary sensory pathways of different modalities. However, the physiological role of dopamine in Nudipleura is only studied in the feeding behaviors of *Pleurbranchaea* (Brown et al., 2023; Norekian et al., 2024).

The revealing systemic integrative (hormonal) roles of dopamine in molluscs is also challenging. In some aspects, work on *Clione* pioneered the field. Injections of dopamine or its metabolic precursor (L-DOPA) suppressed the locomotion/swimming and produced actions opposite to serotonin-induced swimming/feeding behaviors (Sakharov and Kabotyanski, 1986).

Dopamine application on the central ganglia of *Clione limacina* produced differential depolarizing or hyperpolarizing effects on functionally identifiable neurons controlling antagonistic behavioral programs (Norekian, 1990b), perhaps as parts of behavioral states opposite to serotonin-induced behaviors (Sakharov and Kabotyanski, 1986). Dopamine might even directly suppress serotonergic transmission in *Aplysia* (Shozushima et al., 1991). Some dopamine antagonists blocked the spontaneous or evoked inhibitory inputs to the swimming system in *Clione* (Sakharov and Kabotyanski, 1986; Kabotyanski and Sakharov, 1988). Dopamine also activated some neurons involved in the withdrawal (Norekian, 1990b) of *Clione*. As a result, we suggest that dopamine plays a role in the integration of passive defensive pattens with setotonin-dependent behavioral arousal, possibly with the involvement of peripheral neurons as chemoreceptors and modulating mechanoreceptions.

Of note, comparable integrative relationships between DA and serotonin were reported for *Lymnaea* with induction of the overall defensive reaction at higher concentrations with the suppression of locomotion, while smaller concentrations induced well-coordinated respiratory behavior with all its components, including stereotyped inhibition of ciliated and muscular components of locomotion and body orientation, reduction adhesion, mucus secretion and activation of the respiratory central pattern generator (Moroz, 1991). Similar DA-induced behaviors were observed in *Helix pomatia* (Moroz, unpublished observations), and profound inhibition of locomotion of an isolated foot was reported in *H. lucorum* (Pavlova, 2001).

### Conclusions and future directions

Fractions of dopaminergic neurons as related to all other peripheral neurons are apparently different across tissues and species. In rhinophores and other chemosensory systems of *Melibe*, the DA-containing cells represent the majority of neurons, whereas in the wings of pteropods, less than half of peripheral neurons are dopaminergic.

In any scenario, the dopamine and serotonin systems in molluscs are functionally coupled. Nevertheless, obtained data suggest greater phylogenetic plasticity of dopaminergic neurons (in terms of their localization and diversity) compared to serotonergic neurons, which populations are more evolutionary conserved across Euthyneura (Kistler et al., 1985; Longley and Longley, 1986; Satterlie et al., 1995; Marois and Carew, 1997; Sudlow et al., 1998; Dickinson et al., 2000; Delgado et al., 2012).

In most of the studied cases, the action of serotonin is primarily associated with the general arousal and facilitation of synaptic connections by raising the excitability of many neurons in various networks related to enhanced energetic demands such as locomotion and cilia control. By analogy, we view serotonin’s functions and mechanisms as being ‘oil’ in an engine (Moroz, 1991), where serotonergic cells are central interneurons and motoneurons, but there are no serotonergic neurons at the periphery in *Clione* or any other studied euthyneurans (Moroz et al., 1997; Sudlow et al., 1998; Croll et al., 2003). In contrast, dopamine-dependent behavioral and motor patterns are more specific than those induced by serotonin. All known identified dopaminergic central neurons are modulatory interneurons. Still, the CNS receives a majority of dopaminergic inputs from the widespread populations of peripheral neurons, controlling feeding motor patterns (universally) and respiratory behavior in pulmonate molluscs (Moroz, 1990; Moroz, 1991; Moroz and Winlow, 1992a).

Many questions remain regarding to selective forces of neurotransmiters’ evolution (Moroz and Romanova, 2021). Why are dopaminergic cells so abundant and so diverse on the periphery? Why dopamine (vs serotonin)? What are other transmitters within peripheral networks in molluscs? What are the constraints of transmitter-specific synaptic evolution? These and many other fundamental questions require interdisciplinary and comparative studies of these remarkable neural systems and species in Euthyneura and beyond.

## Grant Acknowledgments

This work was supported by the National Science Foundation (IOS-1557923) and the National Institute of Neurological Disorders and Stroke of the National Institutes of Health under Award Number R01NS114491 to L.L.M.

## Acknowledgments

We thank FHL for its excellent microscopy facilities. The National Science Foundation (IOS-1557923) and the National Institutes of Health (R01NS114491) supported this work (to L.L.M.).

## Data Availability Statements

The data that support the findings of this study are available from Dr. T.P. Norekian upon request.

## Conflict of interest

None of the authors has any known or potential conflict of interest, including any financial, personal, or other relationships with other people or organizations within three years of beginning the study that could inappropriately influence, or be perceived to influence their work.

## Role of the authors

All authors had full access to all the data in the study and took responsibility for the integrity of the data and the accuracy of the data analysis. TPN and LLM share authorship equally. Research design: TPN. Acquisition of data: TPN, LLM. Analysis and interpretation of data: TPN, LLM. Drafting of the article: TPN, LLM. Funding: LLM.

## Notes

### Competing Interest Statement

The authors have declared no competing interest.

